# Newly found rat CD103^-^ dendritic cells are the highly immunogenic conventional DC2 subpopulation, corresponding to the known DC subsets in mice and humans

**DOI:** 10.1101/2024.11.12.623329

**Authors:** Yasushi Sawanobori, Tadayuki Ogawa, Hisashi Ueta, Yusuke Kitazawa, Nobuko Tokuda

## Abstract

Dendritic cells (DCs), the primary antigen-presenting cells, have traditionally been identified by CD103 molecules in rats, whereas mouse and human DCs are identified by CD11c molecules. However, this history does not preclude the existence of CD103^-^ DCs in rats. To explore this possibility, we examined MHCII^+^ cells in rat spleen and thymus, identifying a novel population of CD103^-^MHCII^+^CD45R^-^CD172a^+^ cells. These cells are negative for CD103 and B cell marker CD45R, but positive for the type-2 conventional DC (cDC2) marker CD172a. Transcriptomic analyses revealed that they represent a subpopulation of cDC2. Additionally, gene set enrichment analysis predicted enhanced immunogenic activities for this novel population compared to known rat cDC2s. Mixed leukocyte reaction assays confirmed that the rat CD103^-^ cDC2s induce T cell proliferation more effectively than other DC subsets, suggesting enhanced immunogenic potential. In reaggregated thymic organ culture assays, both the rat CD103^-^ and CD103^+^ cDC2 subsets suppressed the total number of generated thymocytes and skewed the differentiation toward CD8 single-positive cells. Comparisons with previously published single-cell RNA-sequencing datasets showed that the rat CD103^-^ cDC2 subset shares markers and GO terms of known mouse and human cDC2 subpopulations: cDC2a, cDC2b, inf-cDC2, and moDC. In contrast, the classic rat CD103^+^ cDC2 subset expresses only cDC2a markers. These findings provide new insights into DC subpopulations, particularly in species other than mice and humans, where much remains to be uncovered.

## Introduction

Dendritic cell (DC) subsets are well-known for their antigen-presenting, cytokine-producing, and stimulatory ability to other immune cells (1). They consist of three main subsets: conventional DC1 (cDC1), cDC2, and plasmacytoid DC. In both thymus and peripheral lymphoid tissues, cDC1 and cDC2 exhibit different localizations and expression of pattern recognition molecules, playing distinct roles with some degree of redundancy. For example, cDC1 subsets mainly induce cytotoxic T lymphocytes and perform cross-presentation, while cDC2 subsets activate CD4^+^ T cells. In the thymus, although eradicating one of the cDC subsets does not simply lead to a failure of central tolerance (2,3), tolerance against certain antigens may depend on specific cDC subsets (4). Moreover, it has been reported that cDC2 subsets play a significant role in capturing and presenting blood-borne antigens (5,6).

Regarding the cDC2 subset, more detailed subpopulations have been identified: cDC2a, cDC2b, inflammatory cDC2 (inf-cDC2), and monocyte-derived DC (moDC) (7). These subpopulations all share the expressions of CD11b, CD11c, CD172a, and MHCII. While cDC2a and cDC2b can be distinguished as ESAM^hi^ and ESAM^lo^, respectively, cDC2b, inf-cDC2, and moDC are easily confused because they share markers such as CCR2, CD64, and Ly6C. Furthermore, cDC2b and moDC exhibit extreme similarities in the expression of CD14, F4/80, CD115, CX3CR1, and other markers. Using tracing models exploiting *Ms4a3* or *Mafb*, it has been suggested that moDCs are derived from monocytes (8,9). The origin of cDC2b remains unclear. Due to the high degree of similarity, cDC2b cells seem to be synonymous with or may at least include moDCs. However, a fate-mapping model suggested that ESAM^lo^ cDC2b can arise from DC precursors (10). On the other hand, inf-cDC2 subpopulations are considered “*bona fide*” DC, characterized by their CD26^hi^CD64^neg-mid^ phenotype (11,12).

As is often the case with other cell types, DC subsets have been extensively studied in mice and humans, and it is well-established that their primary marker is CD11c (13). However, in rats, the monoclonal antibody OX-62, initially established against veiled cells obtained from thoracic ducts of mesenteric lymphadenectomized rats, has been employed as a conventional DC marker (14-19). Later, its epitope was identified as rat CD103 molecules (20). However, this history does not guarantee that anti-CD103 antibodies can detect all rat conventional DC subsets. Confusingly, in mice, CD103 is mainly expressed on cDC1 subsets and is absent from cDC2 subsets, except for intestinal cDC2 (1), while our group has reported that rat CD103^+^ DCs include both XCR1^+^ (a cDC1 marker) cells and CD172a^+^ (a cDC2 marker) cells (21,18).

Our previous study demonstrated the accumulation of CD103^+^XCR1^+^ cDC1 in epithelium-containing areas of rat thymic medullas utilizing immunohistology of serial sections (18). During this investigation, we considered the possibility that some CD103^-^ MHCII^+^ cells might be present in the rat thymus. To confirm this, we performed a flow cytometric analysis on MHCII-enriched thymic cell suspensions and discovered a novel CD103^-^MHCII^+^CD45R^-^CD172a^+^ cell subset. This population is commonly found in the thymus and peripheral lymphoid organs.

In this study, we characterized the CD103^-^MHCII^+^CD45R^-^CD172a^+^ cells from the thymus and spleen, utilizing flow cytometry, transcriptome analysis, and *in vitro* functional assays. The results suggest the CD103^-^MHCII^+^CD45R^-^CD172a^+^ cells are a subpopulation of cDC2. Moreover, comparative analyses revealed that the rat CD103^-^ cDC2 subsets include cDC2b, inf-cDC2, and moDC, corresponding to their mouse and human counterparts and exhibiting high immunogenic potency.

## Materials & Methods

### Animals

Inbred male Lewis rats, 8**–**12 weeks old, were purchased from SLC Co. (Shizuoka, Japan) and reared under specific pathogen-free conditions. The exact ages of the rats used are indicated in the methods or figure legends. As sources of T cells in the mixed leukocyte reactions (MLRs), 8**–**9-week-old male ACI rats were also purchased from SLC Co. Animal handling and care protocols were approved by Dokkyo Medical University’s Regulations for Animal Experiments and adhered to Japanese Governmental Law (No. 105).

### Antibodies

The antibodies used for flow cytometric analysis and cell sorting are listed in *supplementary data and tables.xlsx*. Some antibodies were purified from culture supernatants and conjugated in-house.

### Tissue digestion and cell preparation

Spleens and thymi were injected with an enzyme mixture containing 0.2% collagenase D (Roche Diagnostics, Indianapolis, IN, USA) and 0.01% DNase I (Roche) in Hank’s buffered saline solution (HBSS) with 1.25 mM Ca^2+^, 1.45% bovine serum albumin (BSA), and 2% fetal calf serum (FCS). The tissues were then cut into 1**–**2 mm thick slices and incubated in the enzyme mixture at 37°C for 25 minutes. During incubation, the tissues were pipetted every 5 minutes using truncated 1000 µl pipet tips, followed by Pasteur pipets, and finally, heat-tapered tip Pasteur pipets. After the incubation, EDTA was added to a final concentration of 2.5 mM, and the samples were incubated for an additional 5 minutes. The digested tissues were then filtered using pieces of 50 μm nylon mesh. The resulting suspension was centrifuged and resuspended in PBS(-).

For splenic DC analysis and isolation, cells from digested spleens were resuspended in 15% OptiPrep (Axis-Shield, Oslo, Norway) in PBS(-) and placed in centrifuge tubes. Layers of 11.5% OptiPrep/PBS(-) and PBS(-) were overlaid on the cell suspension. The tubes were centrifuged at 600 *g* for 25 minutes at room temperature with gentle acceleration and braking. Cells between the 15% and 11.5% OptiPrep layers were collected as DC-containing low-density cells and washed with PBS(-).

For thymic DC analysis and isolation, cells from digested thymi were resuspended at 5 × 10□ cells/ml in PBS, and anti-rat MHCII antibody (BioLegend, San Diego, CA, USA) was added at a concentration of 0.5 µg/ml. Cells were incubated at 4°C for 30 minutes and washed with 0.5% BSA and 2 mM EDTA/PBS(-) (MACS buffer). Cells were then resuspended at 20 × 10□ cells/ml in MACS buffer, and 20% anti-mouse IgG microbeads (Miltenyi Biotech, North Rhine-Westphalia, Germany) were added. The cells were incubated at 4°C for 15 minutes, washed twice with MACS buffer, filtered through 50 μm nylon mesh pieces, and resuspended at 1.5 × 10□ cells/ml in MACS buffer. The cells were isolated using the Possel program of autoMACS (Miltenyi Biotech). To increase yield, the negative fraction was subjected to the Possel program again. Later, this isolation method was substituted with a single run of the Posseld2 program.

For thymic and splenic macrophage isolation, cells from digested thymi or spleens were resuspended at 5 × 10□ cells/ml in PBS. Mouse gamma globulin (Jackson ImmunoResearch, West Grove, PA, USA) was added at 20 µg/ml to block nonspecific binding, and the cells were incubated at 4°C for 10 minutes. Biotinylated anti-CD11b/c antibody was added at 0.25 µg/ml, and the cells were incubated at 4°C for 10 minutes, followed by washing with MACS buffer. Cells were then resuspended at 1 × 10^9^ cells/ml in MACS buffer, and 10% streptavidin microbeads (Miltenyi Biotech) were added. After incubation at 4°C for 15 minutes, the cells were washed twice with MACS buffer, filtered with 50 μm nylon mesh pieces, and resuspended at 1.5 × 10□ cells/ml in MACS buffer. The cells were isolated using the Posselds program of autoMACS.

For T cell isolation for MLR assays, cells from digested lymph nodes were resuspended in 15% OptiPrep in a centrifuge tube, and PBS(-) was overlaid on the cell suspension. The tube was centrifuged at 600 *g* for 25 minutes at room temperature. Cells between the 15% OptiPrep and PBS(-) layers were collected as lymphocytes, washed with PBS(-), and resuspended at 3 × 10□ cells/ml in MACS buffer. PE-conjugated anti-rat CD45R antibody (BioLegend) and PE-conjugated anti-CD11b/c antibody (eBioscience) were added at 2.5 µg/ml and 1.5 µg/ml, respectively. The cells were incubated at 4°C for 30 minutes, washed with MACS buffer, and resuspended in MACS buffer at 6 × 10□ cells/ml. Anti-PE microbeads (Miltenyi Biotech) were added at 20%, and the cells were incubated at 4°C for 15 minutes, washed twice with MACS buffer, filtered with 50 μm nylon mesh pieces, and resuspended at 2 × 10□ cells/ml in MACS buffer. The cells were isolated using the Depl025 program of autoMACS. The purity of T cells was confirmed to be over 95%.

### Flow cytometry

Prepared cells were stained following conventional methods. Leucoperm (BIO-RAD, Hercules, CA, USA) was used for CD68 staining. Stained cells were acquired with an Attune NxT flow cytometer (Thermo Fisher Scientific, Waltham, MA, USA). Dead cells were excluded using propidium iodide (PI, Dojindo, Kumamoto, Japan) or 7AAD (IMMUNOSTEP, Salamanca, Spain), following the manufacturer’s protocols. Data were analyzed with FlowJo V10.10.0 (FlowJo LLC, Ashland, OR).

### Cell sorting

The prepared cells were stained following the standard flow cytometry protocol to isolate DC subsets and macrophages. The stained cells were sorted using FACSAria™ III cell sorter (BD, Franklin Lakes, NJ, USA) and collected in HBSS supplemented with 1.25 mM Ca^2+^, 1.45% BSA, and 2% FCS. The sorted cells were then subjected to further experiments. Cytosmears were prepared for morphological observation of the sorted cells, and May-Grünwald-Giemsa staining was performed following the standard procedures.

### RNA microarray

Sorted cells from 8- to 11-week-old Lewis rats were dissolved into ISOGEN (NIPPON GENE, Tokyo, Japan), and RNA was extracted following the manufacturer’s protocol. To ensure DNA removal, the RNA samples were treated with RNase-free DNase I (Takara Bio, Shiga, Japan) following the manufacturer’s protocol. RNA was precipitated with Ethachinmate (NIPPON GENE) and dissolved in sterilized water. All cell types were analyzed using two samples each. Some samples were pooled from multiple animals to ensure sufficient RNA yields. Details of the RNA samples are listed in the *supplementary data and tables.xlsx*.

The microarray was outsourced to Kurabo (Osaka, Japan) and performed using the Clariom™ S Assay, rat (Thermo Fisher Scientific). At least 250 ng of RNA was delivered for outsourcing. RNA integrity numbers were at least 7.9. The data were initially analyzed with the Transcriptome Analysis Console (TAC) v4.0.3 (Thermo Fisher Scientific). Marker genes of mouse and human DC subsets, as well as GSEA gene sets, were applied using the import gene list function. For heatmap generation and comparison with RNA-seq data, data normalized with the TAC were saved as a CSV file and imported into R version 4.4.0.

### Comparison of rat microarray and mouse RNA-seq datasets

The rat microarray expression data, normalized using the TAC, and the mouse bulk or pseudobulk RNA-seq datasets were imported into R. For scRNA-seq data, pseudobulk expression data were generated using the AggregateExpression function with the settings normalization.method = "RC" and scale.factor = 1,000,000. If the expression data were raw counts, they were log-transformed using a base of 2. Genes were converted to rat orthologs using the orthogene package (version 1.9.1). The data were merged using the merge function by row names (gene symbols) and normalized with the removeBatchEffect function from the limma package (version 3.60.2), followed by standardization using the scale function. Heatmaps of gene expressions were generated using the pheatmap package (version 1.0.12).

To assess similarities between rat and mouse/human cell counterparts, population standard deviations (SDs) for each gene between rat cDC1, CD103^+^ cDC2, and CD103^-^ cDC2 were calculated. When the comparison is focused on CD103^+^ cDC2 and CD103^-^ cDC2 subsets or human DC subsets, SDs were calculated only between these specific subsets. Genes were ranked by SDs, and the top 1000 genes were considered differentially expressed genes (DEGs). Pearson correlation coefficients were then calculated between the DEGs of rat and mouse/human subsets, and the results were displayed as heatmaps. In some cases, SDs between Pearson correlation coefficients were calculated to identify corresponding subsets.

To examine marker gene expression within rat transcriptomes, markers of mouse or human subpopulations were extracted from scRNA-seq datasets using the FindAllMarkers function with the settings min.pct = 0.25 and logfc.threshold = 1. The marker genes were converted to rat genes and saved as text files. These text files were loaded into the TAC software as gene lists. Average fold changes and the “differential scores” were calculated among genes in the lists. The differential scores are derived as follows:

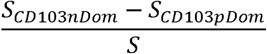

Where S is the total number of genes in the marker gene lists, SCD103nDom is the number of genes dominantly (fold change > 2) expressed in the CD103^-^ subset, and SCD103pDom is the number of genes dominantly (fold change < -2) expressed in the CD103^+^ subset.

### Analyses of single-cell RNA-seq (scRNA-seq) datasets

Downloaded scRNA-seq expression matrices were analyzed using R and the Seurat package (version 5.1.0). Low-quality cells, such as those that were outliers in nFeature_RNA or had a high percentage of mitochondrial RNA, were removed. Clusters and 2D projections were reproduced to match those in the original articles using the reported marker expression. When clustering information or 2D projections were available as metadata tables, we used them directly.

### Gene set enrichment analysis (GSEA)

For microarray datasets, logarithmic expression data were used to calculate log2 fold changes (log2FCs) between subsets. For scRNA-seq datasets, the FindMarkers function in Seurat was used to calculate log2FCs between certain subsets. The resulting log2FC values were compiled into numeric objects labeled with gene names and subjected to GSEA using the gseGO function from the clusterProfiler package (version 4.12.0). Results were visualized as tree plots, running score plots, and enrichment maps using the treeplot, gseaplot2, and emapplot functions, respectively.

To compare results across species, terms with enrichmentScore > 0 were extracted and ranked by their adjusted p-values (p.adjust). This ranking was referred to as *adjRank*, with the term having the lowest p.adjust value assigned *adjRank 1*. Terms within the adjRank range of 1–500 were selected and compared across species. To assess the significance of skewed distributions of *adjRank* values within the shared terms, the Wilcoxon rank-sum test was applied, with the null hypothesis that the *adjRank* values of the shared terms are symmetric about the median of the parent population.

### Mixed leukocyte reaction (MLR)

Isolated T cells from 8–9-week-old ACI rats were stained with CellTrace™ Violet (Invitrogen) following the manufacturer’s protocol. A total of 1 ×10^5^ stained T cells, along with varying numbers of splenic DC subsets isolated from 9– -11-week-old Lewis rats, were seeded into each well of a round-bottom 96-well plate in 200 µl of RPMI-1640 medium (Invitrogen; pre-supplemented with L-glutamine, HEPES, and pyruvate), further supplemented with 1× non-essential amino acids, 50 µM of 2-mercaptoethanol, 1× antibiotic-antimycotic, and 10% FCS. The MLRs were incubated at 37°C for five days. After the incubation, the plates were centrifuged and analyzed following the standard flow cytometry protocol. 7AAD^-^RT1Aa^+^ cells were considered responder T cells, and cells that had divided more than twice were considered dividing.

### Reaggregated thymic organ culture (RTOC)

For RTOCs, fetal thymi at embryonic day 18 were harvested following a previously published method (22). Harvested thymi were kept in ice-cold HBSS supplemented with 1.25 mM Ca^2+^, 1.45% BSA, and 2% FCS during dissection. Pooled fetal thymi were digested in 0.1% collagenase/dispase (Roche) and 0.02% DNase I in HBSS at 37°C for 30 minutes, with pipetting every 10 minutes using 1000 µl and subsequently 200 µl pipet tips. The dissolved thymi were filtered through 50 μm nylon mesh pieces, and the cells were washed twice with PBS. A total of 6.5 × 10^5^ fetal thymic cells and 6.5 × 10^3^ sorted thymic DCs from 8–12-week-old rats per culture were resuspended in DMEM medium (Thermo Fisher Scientific; high glucose, GlutaMAX™ and HEPES supplemented), further supplemented with 1× non-essential amino acids, 50 µM of 2-mercaptoethanol, 1× antibiotic-antimycotic, and 10% FCS, in 1.5 ml microtubes. The tubes were centrifuged at 350 *g* at 4°C for 5 minutes. After centrifugation, the supernatant was completely removed by aspiration and pipetting, and the cell pellets were disaggregated by vortexing. Sterilized 2 cm^2^ cotton pads were placed in 6 cm dishes, and the medium was poured to soak the pads. Hydrophilic polycarbonate membranes with 8.0 µm pore size and 13 mm diameter (Merck Millipore, Burlington, MA, USA) were placed on the soaked pads. The cells were transferred with a 10 µl pipette tip and placed on the membranes in small drops. The dishes were placed in Ziploc plastic containers (SC Johnson, Racine, WI, USA) with 10 ml of sterilized water, and RTOCs were cultured at 37°C for four days. Cultured RTOCs were collected into 1.5 ml microtubes and digested in 100 µl of 0.2% collagenase D and 0.01% DNase I (Roche) in HBSS at 37°C for 30 minutes, with pipetting every 10 minutes using 200 µl pipet tips. After digestion, EDTA was added to a final concentration of 5 mM, and samples were incubated for an additional 5 minutes. The cells were divided into two suspensions for surface marker and intracellular Foxp3 analysis. The cells were analyzed by flow cytometry. 7AAD^-^CD45^+^ cells were considered generated thymocytes, and the ratios of CD4 or CD8 single-positive cells within these cells were calculated. If the number of generated thymocytes was fewer than 3500, the RTOC was considered defective and excluded from analysis. Data from independent experiments were pooled and presented as relative values to the control (23).

### Statistical analyses

For *in vitro* assays, the Student’s t-test was used to assess significance. The source data for each figure are included in the *supplementary data and tables.xlsx*.

## Result

### Both the rat spleen and thymus contain CD103^-^MHCII^+^CD45R^-^CD172a^+^ cells

To survey all antigen-presenting cells in the rat thymus, we used magnetic cell sorting targeting rat MHCII molecules, achieving an approximately 80% concentration of MHCII^+^ cells (Fig. S1A). For the spleen, we applied a density gradient to enrich low-density cells, as the high number of B cells in the MHCII-magnetically sorted fraction complicated analysis. Due to the strong autofluorescence of PI^dull^ cells in the splenic low-density fraction, we excluded these cells from further analyses (Fig. S1B). Both splenic and thymic cells contained CD103^+^ conventional rat DCs, which included CD172a^-^ (and XCR1^+^, discussed later) cDC1 and CD172a^+^ cDC2 subsets (Fig. 1A). Additionally, within the CD103^-^MHCII^+^ fractions, some CD45R^-^CD172a^+^ cells were observed. May-Grünwald-Giemsa staining revealed that cDC1 cells were relatively large and displayed delicate protrusions in both the thymus and spleen (Fig. 1B). In contrast, some cDC2s and CD103^-^MHCII^+^CD45R^-^CD172a^+^ cells were small and exhibited pseudopod-like protrusions. Regarding cell abundance, cDC2 cells were the most common in the spleen, followed by cDC1 cells, while CD103^-^MHCII^+^CD45R^-^ CD172a^+^ cells were the least abundant (Fig. 1C). In the thymus, cDC1 was the most prevalent population, with cDC2 and CD103^-^MHCII^+^CD45R^-^CD172a^+^ cells being less abundant, with no significant difference between the latter two.

**Figure 1.**
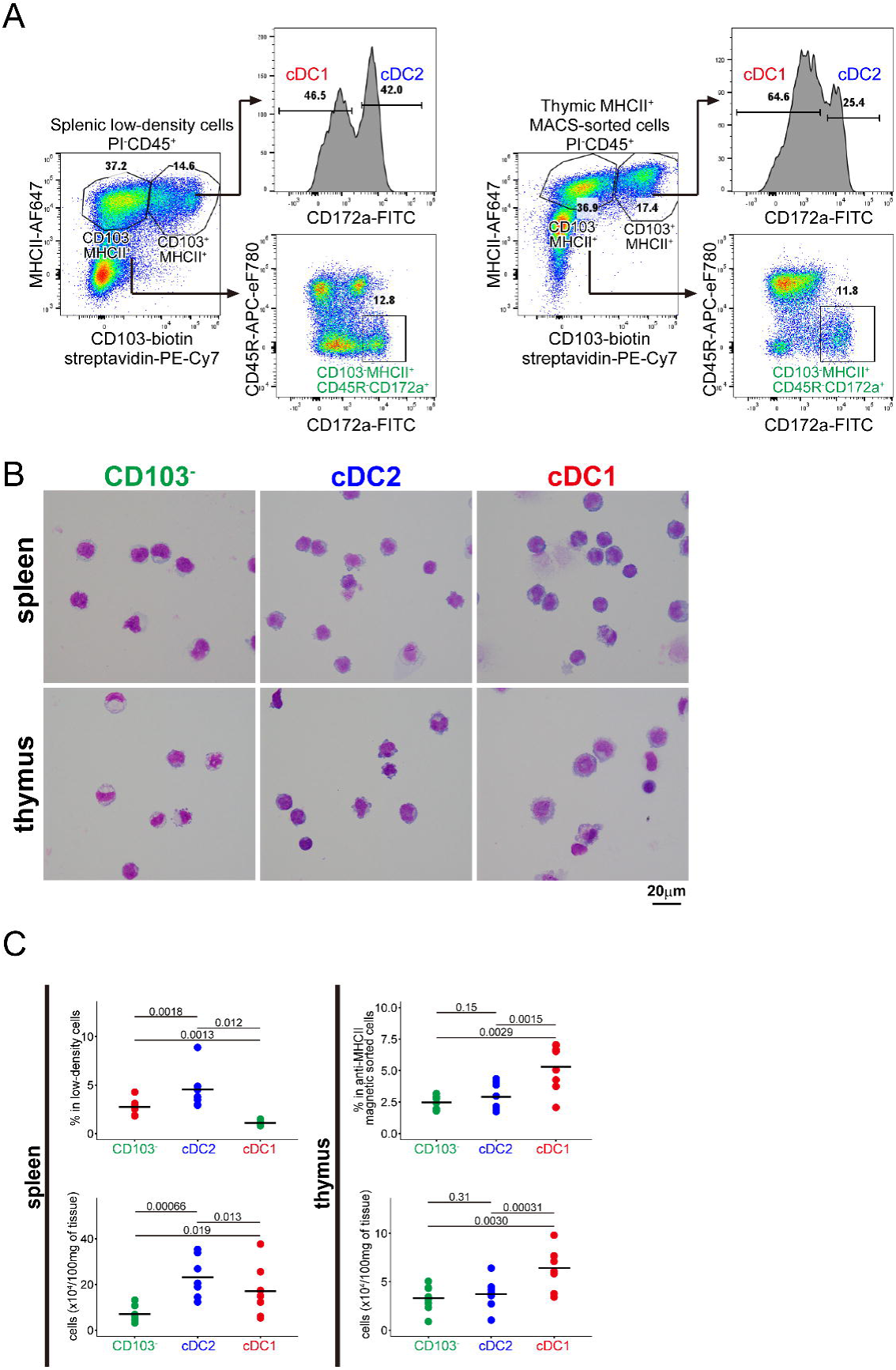
CD103^-^MHCII^+^CD45R^-^CD172a^+^ cells are found in rat lymphoid tissues. (A) Digested rat spleens and thymi were subjected to flow cytometric analysis. In addition to CD103^+^ cDCs, CD45R^-^CD172a^+^ cells were identified within the CD103^-^ MHCII^+^ subsets. Representative plots from an 8-week-old Lewis rat are shown. (B) Images of May-Grünwald-Giemsa-stained sorted subsets. The cells were isolated from 10-week-old Lewis rats. (C) Percentages and absolute numbers of the three subsets. P-values from Student’s t-tests are displayed.

Flow cytometric analysis was performed to investigate surface marker expression (Fig. S1C). As anticipated, the cDC1 marker XCR1 was exclusively expressed on cDC1 cells. CD26, which helps to distinguish moDCs from *bona fide* DCs through low expression (11,12), was also low on CD103^-^MHCII^+^CD45R^-^CD172a^+^ cells. CD161, known as a marker for both NK cells and DCs in rats (24,25), was expressed on all three cell types in both the spleen and thymus. However, some splenic CD103^-^MHCII^+^CD45R^-^CD172a^+^ cells lacked CD161 expression, suggesting the need for further gating for the isolation. Unlike in mice, where CD11b is considered a cDC2 marker (1,13), all rat DC subsets partially expressed CD11b and CD11c. As noted in our previous reports (21,18), thymic cDC1 and cDC2 displayed CD4^dull^CD8a^dull^ and CD4^+^CD8a^dull^ profiles, respectively, while splenic cDC1 and cDC2 were CD4^dull^CD8a^-^ and CD4^+^CD8a^-^. The CD103^-^CD45R^-^ CD172a^+^MHCII^+^ cells exhibited a cDC2-like pattern of CD4 and CD8a expression. Costimulatory molecules CD80 and CD86 showed similar expression patterns across all subsets, except for higher CD80 expression on splenic cDC2. Although CD68, CD163, and CD169 are traditionally used as macrophage markers in rats (26,27), CD68 was expressed in all three cell subsets, whereas CD163 and CD169 were found in only a small fraction. To confirm that CD103^-^MHCII^+^CD45R^-^CD172a^+^ cells are exclusively CD172a^+^XCR1^-^, we gated the CD103^-^MHCII^+^CD45R^-^CD161^+^ cells (Fig. S1D). These cells were indeed CD172a^+^XCR1^-^ in both the spleen and thymus, while CD103^+^MHCII^+^ cDCs consisted of XCR1^+^ cDC1 and CD172a^+^ cDC2.

### The transcriptome analysis revealed that CD103^-^MHCII^+^CD45R^-^CD172a^+^ cells represent novel subsets of cDC2

To compare their transcriptomes with microarray analyses, cDC1, cDC2, and CD103^-^ MHCII^+^CD45R^-^CD172a^+^ cells were purified using gating strategies similar to those in Fig. 1A and Fig. S1A–B (Fig. S2A, B). For splenic DCs, CD161 gating was further applied to ensure the exclusion of non-DC cells (Fig. S2A), given the CD161 expression on splenic CD103^-^MHCII^+^CD45R^-^CD172a^+^ cells (Fig. S1C).

We also aimed to prepare macrophages for comparison. Two approaches were considered for macrophage isolation: 1) expression of CD68, CD163, and CD169 (27,28), and 2) use of the anti-rat macrophage antibody HIS36 (29-32). Since all DC subsets in this study expressed CD68 (Fig. S1C), we excluded this molecule as a macrophage marker. Among thymic and splenic low-density cells, CD163 was expressed on a small percentage of both MHCII^+^ and MHCII^-^ cells, while CD169 expression was scarce (Fig. S2C). Therefore, CD163 was chosen as the primary macrophage marker. As the epitope of HIS36 has not yet been determined, we examined the relationship between CD163 expression and HIS36 staining (Fig. S2D). Anti-CD163 antibody (ED2) and HIS36 showed an identical staining pattern, suggesting that HIS36 targets CD163. Consequently, we used HIS36^+^ (presumably CD163^+^) cells in CD11b/c-enriched samples as standard macrophages (Fig. S2E).

All microarray runs were performed in duplicate, with some RNA samples derived from multiple animals. The animals used for the RNA samples are summarized in the *supplementary data and tables.xlsx*.

We first used principal component analysis (PCA) to summarize differences between the transcriptomes (Fig. 2A). The same subsets from different organs were positioned close to each other on the PCA plot, suggesting that intra-organ differences between cell subsets are greater than inter-organ differences within the same cell subsets. This finding was confirmed by sources of variation analyses (Fig. 2B). The distances between cell subsets on the PCA charts corresponded to the number of differentially expressed genes (Fig. 2C).

**Figure 2.**
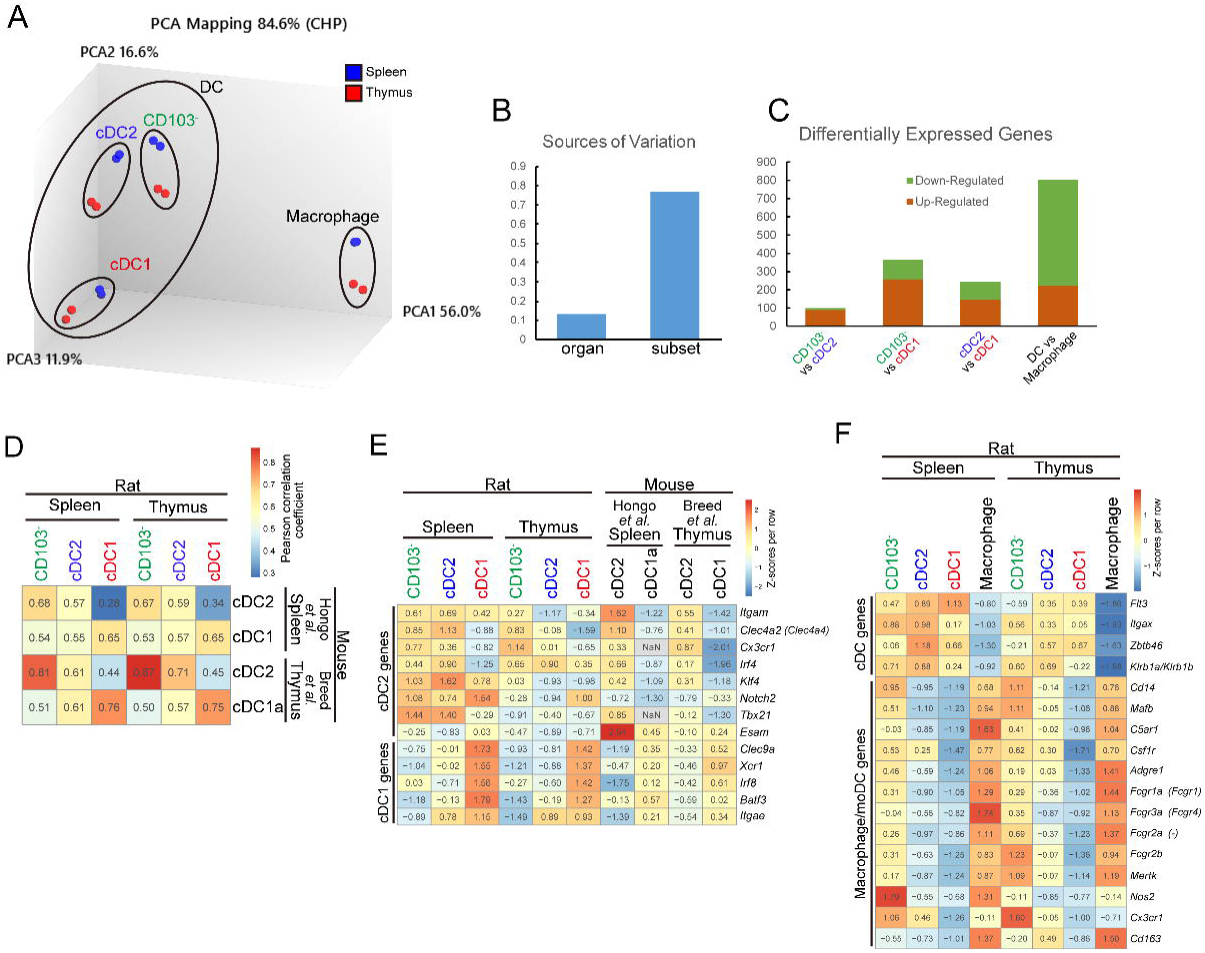
Transcriptomic analyses reveal that CD103^-^MHCII^+^CD45R^-^CD172a^+^ cells are a highly potent subpopulation of cDC2. (A) PCA of cDC1, classic cDC2, CD103^-^MHCII^+^CD45R^-^CD172a^+^ subsets, and HIS36^+^ macrophages. Each subset was derived from both spleens and thymi and includes two samples. Cells were isolated from 8–11-week-old Lewis rats. (B) Sources of variation values calculated by the TAC. (C) Number of differentially expressed genes between indicated combinations. Genes with a p-value < 0.05 and fold change > 10 or < -10 were considered differentially expressed. (D) Pearson correlation coefficients between rat and mouse cDC subsets displayed as a heatmap. The mouse splenic cDC2 transcriptome was calculated from CD301b^+^ and CD301b^-^ cDC2 subpopulations, which comprise 46% and 54% of the parent subset, respectively. (E) Expression of cDC2 and cDC1 markers in rat and mouse cDC subsets. For genes converted from mouse to rat, the original mouse orthologs are shown in parentheses. Data are normalized per row, with Z-scores representing relative expression levels. (F) Expression of DC and macrophage/moDC markers in rat DC and macrophage subsets. Mouse orthologs are indicated in parentheses. Data are normalized per row, with Z-scores representing relative expression levels.

To explore similarities and differences between rat cDC subsets and their mouse counterparts, we compared the rat microarray datasets with mouse bulk RNA-seq datasets (GSE198789 and GSE181475) (33,34). For splenic cDC1, we used the transcriptome of CD8^hi^ cDC1a, as the CD8^lo^ cDC1b subset showed poor similarity to any rat cDCs. Since the original study divided mouse thymic cDC2s into CD301b^+^ and CD301b^-^ fractions, we generated a composite transcriptome for total cDC2 based on the ratio reported in the original paper. We calculated Pearson correlation coefficients to assess the similarities between rat and mouse cDC subsets (Fig. 2D). The Pearson coefficients for the rat CD103^-^MHCII^+^CD45R^-^CD172a^+^ subsets and mouse cDC2 subsets were higher than those for the rat classical cDC2 subsets and mouse cDC2 subsets, suggesting that the rat CD103^-^MHCII^+^CD45R^-^CD172a^+^ subsets are more similar to mouse cDC2s. The rat cDC1 subsets from both the spleen and thymus showed high correlations with mouse cDC1 subsets.

Based on the result of the similarity analysis, we examined the expression of cDC2 and cDC1 markers in the three rat cDC subsets. *Itgam* (CD11b) was expressed in all rat splenic cDC subsets, whereas in mice, it was restricted to cDC2 subsets (Fig. 2E). Other cDC2 markers, such as *Clec4a2* (the ortholog of mouse *Clec4a4*), *Cx3cr1*, *Irf4*, *Klf4*, *Notch2*, *Tbx21*, and *Esam* did not show notable differences between the rat CD103^-^MHCII^+^CD45R^-^CD172a^+^ subsets and cDC2 subsets, particularly in the spleen. Since *Notch2*, *Esam*, and *Tbx21* are reported to be expressed in or required for differentiation of cDC2a, while Klf4 is essential for cDC2b (7), these results suggest the need for further characterization. The cDC1 markers *Clec9a*, *Xcr1*, *Irf8*, and *Batf3* were specifically expressed in cDC1 subsets in both rat and mouse DCs. As expected, *Itgae* (CD103) was expressed in both CD103^+^ cDC2 and cDC1 subsets in rats, while in mice, it was specific to cDC1 subsets.

To confirm that the three rat DC subsets are indeed DCs, we examined the expression of dendritic cell markers. *Flt3*, *Itgax* (CD11c), *Zbtb46*, and *Klrb1a/Klrb1b* (CD161a/CD161b) were moderately to highly expressed in all rat DC subsets but showed low expression in macrophages (Fig. 2F). Considering the potential presence of an moDC subset in rat DC subsets, we also examined the expression of macrophage/moDC marker genes. Many macrophage/moDC markers, such as *Cd14*, *Mafb*, *C5ar1* (CD88), and *Csf1r* (CD115), were expressed both in the CD103^-^ MHCII^+^CD45R^-^CD172a^+^ subsets and the macrophages. *Fcgr1a* (CD64), *Fcgr3a* (rat CD16), the ortholog of MAR-1 epitope *Fcgr4* (35), as well as *Fcgr2a* and *Fcgr2b* (rat CD32), were also expressed in both the CD103^-^ cDC2 and the macrophages. *Nos2* was strongly expressed in the splenic CD103^+^ cDC2 subset and macrophages but not in the thymic counterparts. As expected, *Cd163* was exclusively expressed in the macrophages. Based on these results, we designated the rat CD103^-^MHCII^+^CD45R^-^CD172a^+^ subsets as CD103^-^ cDC2 and the classical cDC2 subsets as CD103^+^ cDC2, respectively.

### Gene set enrichment analysis revealed that the CD103^-^ cDC2 subsets are highly immunologically potent cells

We performed gene set enrichment analysis (GSEA) to examine the biological processes of the CD103^-^ and CD103^+^ cDC2 subsets, analyzing the splenic and thymic subsets separately. The full GSEA results are available in supplementary tables 1–4. To summarize these findings, we used the treeplot function to group the 30 terms with the lowest adjusted p-values (p.adjust) (Fig. 3A, C). In both the spleen and thymus, the CD103^-^ cDC2 subsets exhibited groups indicating enhanced immune responses (e.g., “defense interleukin-1 II interferon” and “lipopolysaccharide molecule bacterial origin”), with p.adjust values reaching saturation (1e-10), suggesting strong differences compared to the CD103^+^ cDC2 subsets. Conversely, the CD103^+^ cDC2 subsets showed groups primarily associated with the cell cycle (e.g., “cycle G2/M phase transition” and “spindle microtubule cytoskeleton involved”), though with relatively higher p.adjust values, ranging from 5.0e-06 to 3e-05.

**Figure 3.**
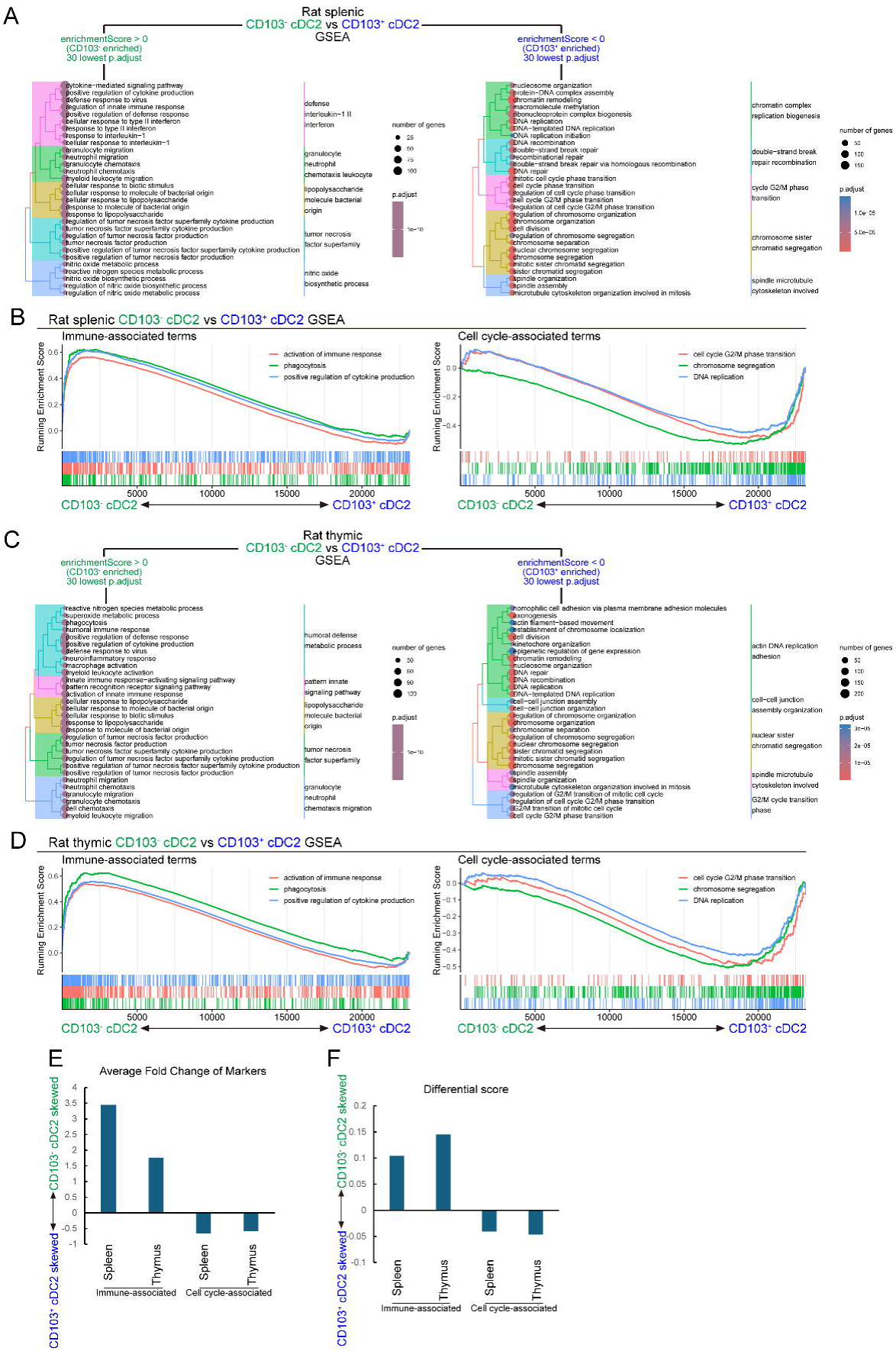
Gene set enrichment analyses of the rat CD103^-^ and CD103^+^ cDC2 subsets. GSEAs were performed between rat splenic CD103^-^ cDC2 and CD103^+^ cDC2 subsets (A, B) and thymic CD103^-^ cDC2 and CD103^+^ cDC2 subsets (C, D). The results are divided into CD103^-^ cDC2-enriched and CD103^+^ cDC2-enriched terms and displayed as tree plots (A, C). Representative gene sets for immune-associated terms (GO:0006909, GO:0002253, GO:0001819) and cell cycle-associated terms (GO:0006260, GO:0007059, GO:0044839) terms are shown as running plots (B, D). (E, F) Marker gene sets were extracted from immune and cell cycle-associated terms. The average fold changes (E) and differential scores (F) between the rat CD103^-^ and CD103^+^ cDC2 subsets were calculated within each marker set.

To visualize gene set enrichments, we selected representative terms for the immune-associated and cell cycle-associated gene sets and illustrated them in running score plots (Fig. 3B, D). In both the thymus and spleen, immune-associated terms (“activation of immune response”, “phagocytosis”, and “positive regulation of cytokine production”) were enriched in the CD103^-^ cDC2 subsets. Meanwhile, cell cycle-associated terms (“cell cycle G2/M phase transition”, “chromosome segregation”, and “DNA replication”) were enriched in the CD103^+^ cDC2 subsets.

To further validate the GSEA results, we extracted genes from the immune-associated and cell cycle-associated terms and applied them to the rat transcriptomes (Fig. S3). Both the average fold changes (Fig. 3E) and the “differential scores” (see the Materials & Methods, Fig. 3F) confirmed that immune-associated genes were enriched in the CD103^-^ cDC2 subsets, while cell cycle-associated genes were enriched in the CD103^+^ cDC2 subsets. However, the absolute values for the cell cycle-associated gene set were smaller than those of the immune-associated genes (0.59-0.65 vs. 1.8-3.4 in average fold changes, 0.041-0.046 vs. 0.10-0.14 in differential scores), consistent with the lower p.adjust values in the GSEAs.

### The characteristics found in the transcriptome analyses were confirmed in in vitro examinations

To validate the enhanced immunologic potency of the CD103^-^ cDC2 subsets *in vitro*, we performed mixed leukocyte reactions (MLRs) and reaggregated thymic organ cultures (RTOCs) for the splenic and thymic CD103^-^ cDC2 subsets, respectively.

For the MLR assays, splenic cDC fractions were mixed with MHC-unmatched T cells (Fig. S4A). The splenic CD103^-^ cDC2 subset induced significantly greater T cell proliferation than the other two cDC subpopulations, as measured by both the percentages and numbers of proliferating cells (Fig. 4A). Notably, we did not separate CD4^+^ and CD8^+^ T cells for the MLR assays, as it was uncertain which T cells would proliferate primarily, and there might be interactions between these subsets via humoral factors. As a result, all three cDC subsets primarily induced CD4^+^ T cell proliferation (Fig. S4B), whereas only the cDC1 subset induced proliferation in a small portion of TCR^hi^CD8^+^ T cells (data not shown).

**Figure 4.**
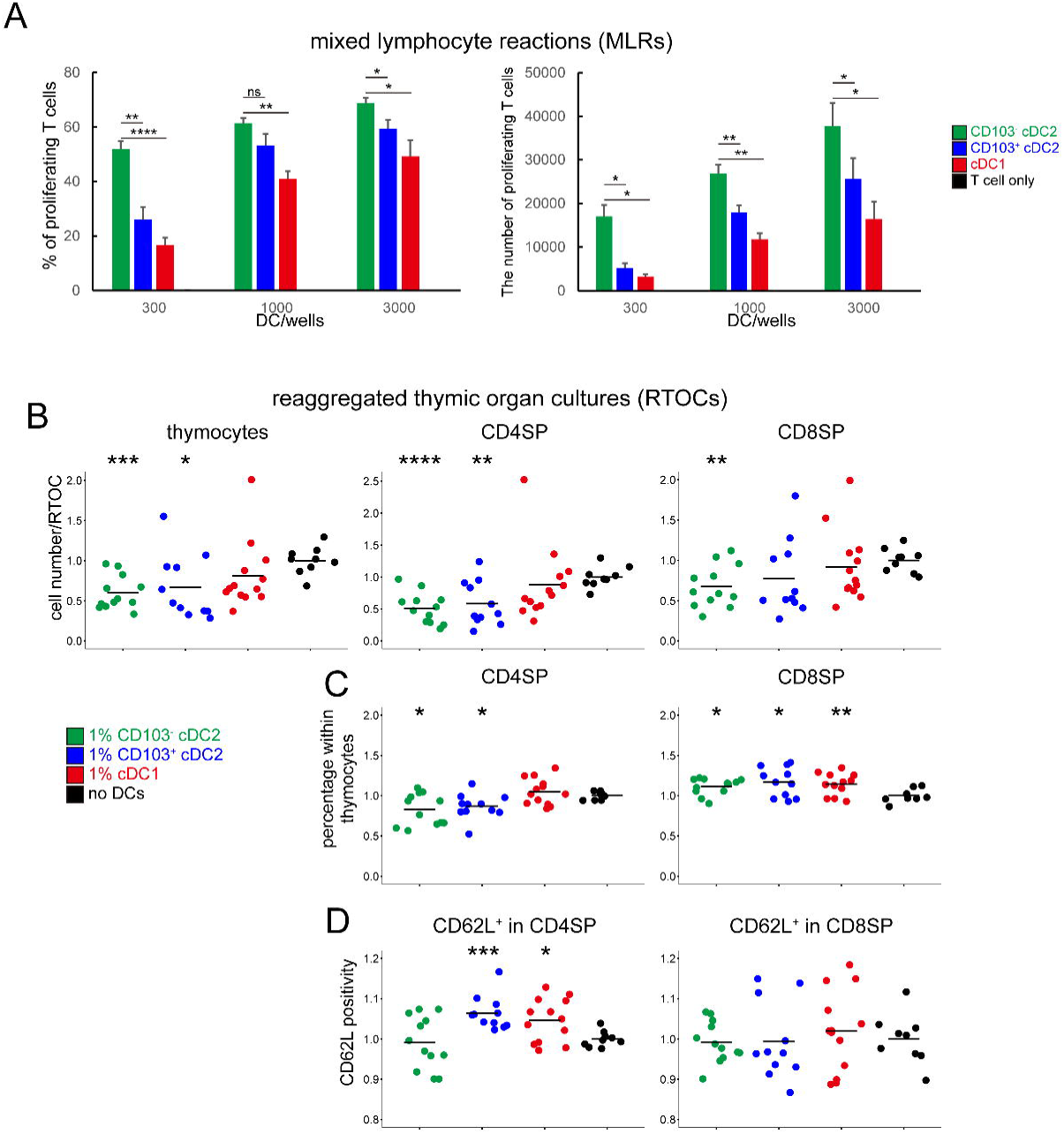
In vitro analyses reveal that CD103^-^ cDC2s are highly potent in T cell priming and negative selection. (A) MLRs were performed using Lewis DCs and ACI T cells. The percentage of proliferating (CellTrace-negative fraction) T cells and absolute numbers of T cells per well are shown. The MLRs were performed in triplicates. Representative data from three independent experiments are shown. *: p < 0.05. (B–D) RTOCs were performed with fetal thymic cells mixed with 1% of thymic DCs sorted from adult rats. The total number of thymocytes (7AAD^-^CD45^+^ cells), CD4SP, and CD8SP subsets (B), as well as the percentages of CD4SP and CD8SP (C) and CD62L^+^ cells (D), are displayed. Data from three independent experiments were pooled and shown as relative values against the controls. *: p < 0.05, **: p < 0.005, ***: p < 0.0005, ****: p < 0.00005.

In the RTOC experiments, a four-day culture of E18 rat fetal thymic cells effectively facilitated reaggregation and generated single-positive thymocytes (Fig. S4C). Once the experimental protocol was optimized, sorted thymic cDC fractions were added to RTOCs at a 1% ratio. Both the CD103^-^ cDC2 and CD103^+^ cDC2 subsets suppressed the overall number of thymocytes within each RTOC, particularly reducing the counts of CD4 single-positive (CD4SP) cells in the cultures (Fig. 4B), suggesting enhanced negative selection (23). The percentages of CD4SP cells within the thymocyte population decreased in the CD103^-^ cDC2- and CD103^+^ cDC2-added RTOCs, while the percentages of the CD8SP cells increased (Fig. 4C). Since CD62L is expressed on terminally differentiated thymocytes (36,37), we analyzed CD62L expression to assess thymocyte maturity (Fig. 4D). The cDC1 and CD103^+^ cDC2 subsets, but not the CD103^-^ cDC2 subset, increased CD62L positivity in the CD4SP population. Measurements of regulatory T cells within the CD4SP population were also attempted, but the data were not statistically significant and, therefore, are not shown.

### The rat CD103^-^ cDC2 subsets show similarities to mouse cDC2a, cDC2b, inf-cDC2, moDC, and macrophages, while the CD103^+^ cDC2 subsets are similar only to cDC2a

To determine which mouse cDC2 subpopulation the rat CD103^-^ and CD103^+^ cDC2 subsets correspond to, we downloaded a gene expression matrix from a single-cell RNA-seq study of mouse splenic CD11c^+^MHCII^+^ cells (38). The accession number is GSM5941846. Clusters were reproduced following the distributed metadata (Fig. S5A). Gene expression data were extracted from the scRNA-seq clusters for the similarity analyses, and Pearson correlation coefficients were calculated between the rat DC transcriptomes and the mouse expression data.

In line with the previous comparison using mouse bulk RNA-seq datasets (Fig. 2D), the rat splenic CD103^-^ cDC2 subset exhibited higher correlations with mouse cDC2 subsets than the CD103^+^ cDC2 subset. Specifically, the CD103^-^ cDC2 subset showed especially high correlation coefficients with mature cDC2a, cDC2b, and monocytes, whereas the CD103^+^ cDC2 subset only showed similarities with cDC2a subpopulations (Fig. 5A). To further assess the differences in the correlation coefficients between the rat CD103^-^ and CD103^+^ cDC2 subsets, we calculated population standard deviations (SDs) between the coefficients of the CD103^-^ and CD103^+^ cDC2 subsets. SDs were notably large for correlations with cDC2b subpopulations and monocytes.

**Figure 5.**
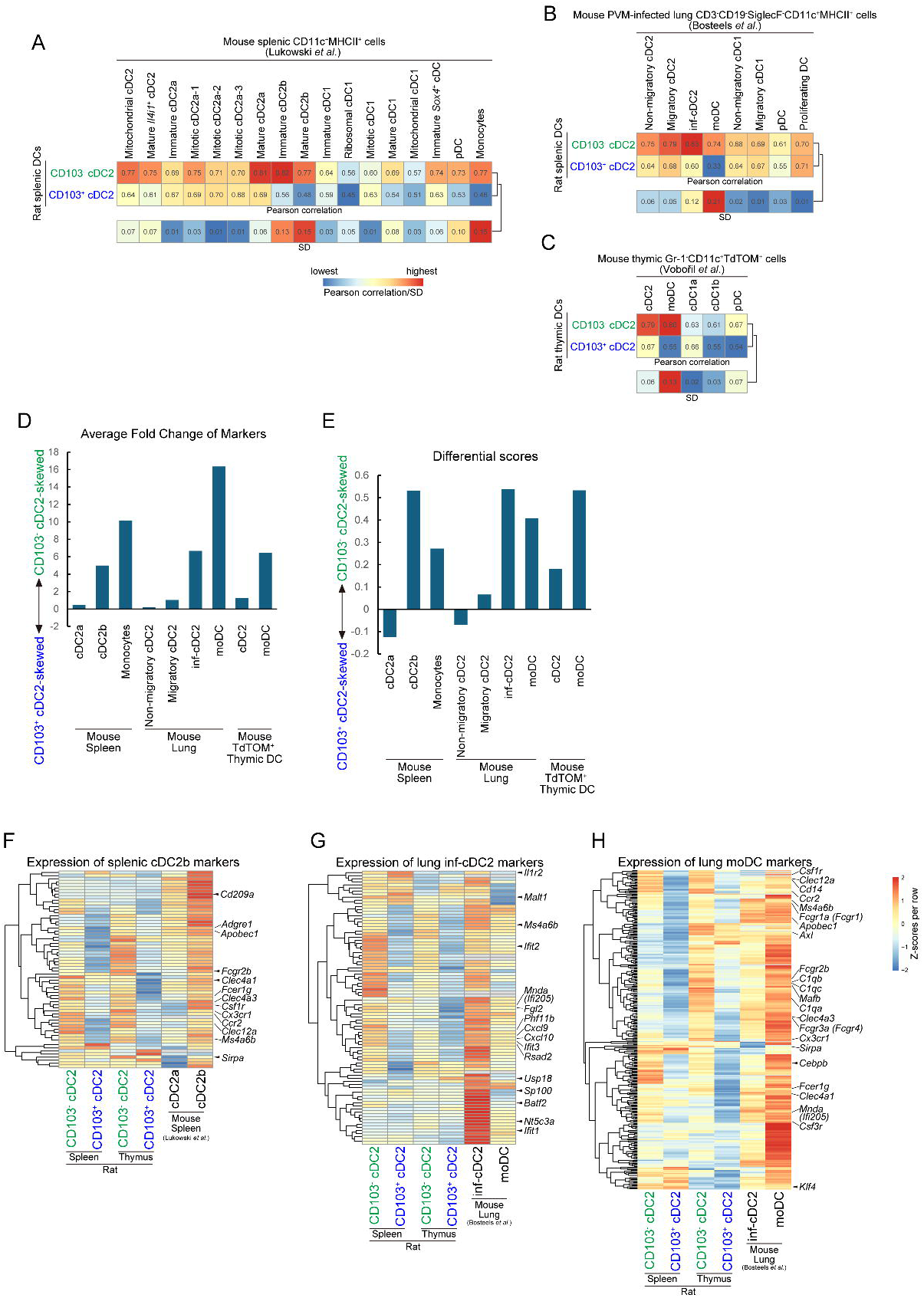
Similarity analyses between rat cDC subsets and mouse cDC subpopulations. (A–C) Pearson correlation coefficients between rat CD103^-^ and CD103^+^ cDC2 subsets and mouse DC subpopulations from different tissues: splenic DCs (A), PVM-infected lung DCs (B), and thymic TdTOMATO-incorporating DCs (C). SDs were also calculated between correlations of rat CD103^-^ and CD103^+^ cDC2 subsets. (D, E) Marker gene sets were generated from mouse scRNA-seq datasets. Mouse subpopulations within the cDC2a and cDC2b subsets were merged to generate marker sets. The average fold changes (D) and differential scores (E) between the rat CD103^-^ and CD103^+^ cDC2 subsets were calculated within each marker set. (F–H) Expression of mouse cDC2b markers (F), mouse lung inf-cDC2 markers (G), and lung moDC markers (H) in the rat CD103^-^ and CD103^+^ cDC2 subsets, displayed as heatmaps. The indicated genes are manually curated. Original mouse orthologs are shown in parentheses for genes converted from mouse to rat. Data are normalized per row, with Z-scores representing relative expression levels.

At this point, we considered that the observed similarity of the rat CD103^-^ cDC2 subset to the monocyte subset might indicate that the rat CD103^-^ cDC2 cells possess an moDC gene signature. To explore this possibility, we utilized an scRNA-seq dataset of pneumonia virus of mice (PVM)-infected mouse lung DCs, which carefully distinguishes cDC2, inf-cDC2, and moDC subsets (12). This dataset was downloaded from http://bioit2.irc.ugent.be/cdc2/download.php, and clusters were reproduced following the distributed metadata (Fig. S5B). As expected, the rat splenic CD103^-^ cDC2 subset displayed a high correlation with the mouse lung moDC subset, while the rat splenic CD103^+^ cDC2 subset exhibited a low correlation (Fig. 5B). Additionally, the rat splenic CD103^-^ cDC2 subset showed a high correlation with inf-cDC2.

Finally, to confirm the high correlation between rat CD103^-^ cDC2 and mouse moDC in the thymus, we referred to an scRNA-seq dataset of thymic DCs incorporating medullary thymic epithelial cell (TEC)-derived TdTOMATO protein (39). The accession number is E-MTAB-8028. The clusters were reproduced and visualized as similarly as possible to the original research (Fig. S5C). Rat thymic CD103^-^ cDC2 strongly correlated with mouse thymic moDC, while the rat thymic CD103^+^ cDC2 exhibited a much lower correlation (Fig. 5C).

### Markers of mouse cDC2b, inf-cDC2, and moDC are predominantly expressed in the CD103^-^ cDC2 subsets

To examine the expression of mouse subpopulation marker genes in the rat transcriptomes, we extracted marker genes from the mouse scRNA-seq datasets using the FindAllMarkers function and applied them to the rat microarray datasets (Fig. S5D–F). Marker genes of mouse splenic cDC2b, splenic monocytes, lung inf-cDC2, lung moDC, and thymic moDC displayed higher expression in the rat CD103^-^ cDC2 subsets. These marker genes are listed in supplementary table 5. To assess the skewed expression toward one of the cDC2 subsets, we calculated the average fold changes (Fig. 5D) and “differential scores” (see Materials & Methods) (Fig. 5E). Both the average fold changes and differential scores indicated that markers for splenic cDC2b, splenic monocytes, lung inf-cDC2, lung moDC, and thymic moDC were predominantly expressed in the CD103^-^ cDC2 subsets.

Next, we analyzed the similarities between marker gene sets. Jaccard similarity coefficients were calculated between the marker sets of mouse subpopulations predominantly expressed in the rat CD103^-^ cDC2 subsets (Fig. S5G). Lung inf-cDC2 markers showed notably low similarities to those of cDC2b, monocyte, and lung moDC subpopulations, indicating that the inf-cDC2 possesses unique marker genes distinct from these other subsets. Thymic moDC subset markers showed similarities with splenic cDC2b, lung inf-cDC2, and lung moDC subsets. These results suggest two key findings: 1) there are two marker set types—the cDC2b/moDC-type, which includes splenic cDC2b, splenic monocytes, and lung moDC markers, and the inf-cDC2 type, which includes only lung inf-cDC2; 2) thymic moDCs have both types of marker sets.

We further examined marker gene expressions using heatmaps, selecting the splenic cDC2b marker, lung inf-cDC2 marker, and lung moDC marker sets as representatives (Fig. 5F–H). The majority of the splenic cDC2b, lung inf-cDC2, and lung moDC markers were predominantly expressed in the CD103^-^ cDC2 subsets compared to the CD103^+^ cDC2 subsets, in both the thymus and spleen. We also curated some genes from related research (12,7) and reviewed their expression on the heatmaps. As expected, the cDC2b markers and moDC markers shared many common genes, such as *Clec4a1*, *Clec4a3*, *Clec12a*, *Fcgr2b*, *Fcer1g*, *Ccr2*, and *Csf1r*. On the other hand, within inf-cDC2 markers, interferon-activated antiviral genes such as *Ifit2*, *Ifit3*, and *Rsad2*, along with chemokines *Cxcl9* and *Cxcl10* and the differentiation marker *Mnda* (*Ifi205*), were predominantly expressed in the rat CD103^-^ cDC2 subsets. In the moDC-specific markers, CD64 (*Fcgr1a*, *Fcgr1* in mouse), the MAR-1 epitope *Fcgr3a* (*Fcgr4* in mouse) (35), and the monocyte lineage marker *Mafb* (9) were also detected.

### The rat splenic CD103^-^ cDC2 subset includes counterparts of human CLEC10A^+^ cDC2 and CCR7^+^ cDC2 subpopulations

To compare the rat cDC2 transcriptomes to human cDC2 subpopulations, we downloaded scRNA-seq data from a published study on human splenic DCs (40), accession number GSM4504590. Clusters and the tSNE projection were reproduced based on the distributed metadata (Fig. S6A). For the comparison, Pearson correlation coefficients were calculated as shown in Fig. 5A–C (Fig. 6A). The splenic rat CD103^-^ cDC2 subsets showed similarities to all human cDC2 subpopulations, whereas the correlation coefficients for the CD103^+^ cDC2 subset were notably low when compared to human CLEC10A^+^ cDC2b and CCR7^+^ cDC2 subpopulations, resulting in higher SDs. Marker genes were identified and applied to the rat transcriptomes (Fig. S6B), and average fold changes and differential scores were calculated (Fig. 6B, C). The marker genes are listed in the supplementary table 5. As expected from the Pearson correlation analysis, the average fold changes and differential scores were relatively high for markers of the human CLEC10A^+^ cDC2b and CCR7^+^ cDC2 subpopulations. However, their maximum values—2.9 and 0.34, respectively—were much lower than those observed in the comparisons with mouse data (Fig. 5D, E), suggesting that the similarities between rats and humans are lower than those within rodent species.

**Figure 6.**
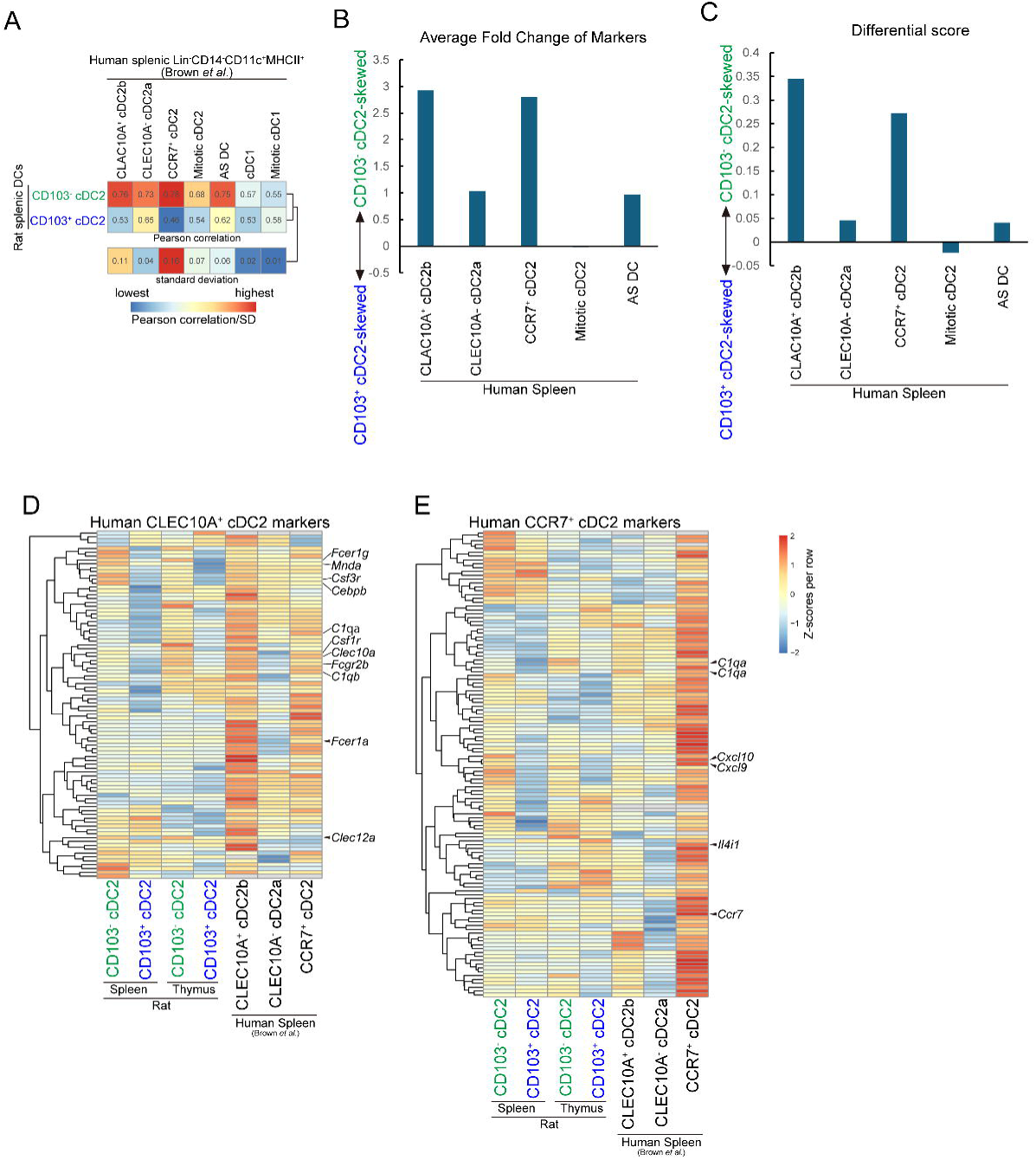
Similarity analyses between rat cDC subsets and human cDC subpopulations. (A) Pearson correlation coefficients between the rat CD103^-^ and CD103^+^ cDC2 subsets and human splenic DC subpopulations. (B, C) Marker gene sets were generated from human scRNA-seq dataset. The average fold changes (B) and differential scores (C) between the rat CD103^-^ and CD103^+^ cDC2 subsets were calculated within each marker set. (E, E) Expression of human splenic CLEC10A^+^ cDC2 markers (E) and CCR7^+^ cDC2 lung moDC markers (F) in the rat CD103^-^ and CD103^+^ cDC2 subsets, displayed as heatmaps. The indicated genes are manually curated. Data are normalized per row, with Z-scores representing relative expression levels.

To provide an overview of the expression of CLEC10A^+^ cDC2b and CCR7^+^ cDC2 markers in the rat cDC2 subsets, we visualized the data using heatmaps (Fig. 6D, E). Genes reported as markers for cDC2b and moDC — *Fcgr1a*, *Fcgr2b*, *Mnda*, *Cebpb*, *Csf1r*, *Csf3r*, *Clec10a*, and *Clec12a* — were among the CLEC10A^+^ cDC2b markers and were predominantly expressed in the rat CD103^-^ cDC2 subsets. For CCR7^+^ cDC2 markers, complement genes *C1qa* and *C1qb*, along with chemokines *Cxcl9* and *Cxcl10,* showed predominant expression in the CD103^-^ cDC2 subsets in both the rat spleen and thymus. However, *Ccr7* and *Il4i1* expression was concentrated in the CD103^+^ cDC2 subset in the rat thymus.

### Commonly enriched terms suggest the significance of CD103^-^ cDC2 equivalent subsets in mice and humans

To identify commonly enriched biological processes, we further performed GSEAs on CD103^-^ equivalents in the mouse and human datasets in addition to the rat datasets analyzed earlier (Fig. 3) and compared the results across species. These GSEAs were conducted on splenic and thymic subsets (Fig. 7A, C). From each GSEA result, the 500 terms with positive enrichmentScore (enriched in the CD103^-^ cDC2 equivalent subsets) and the lowest p.adjust values were extracted for comparison.

**Figure 7.**
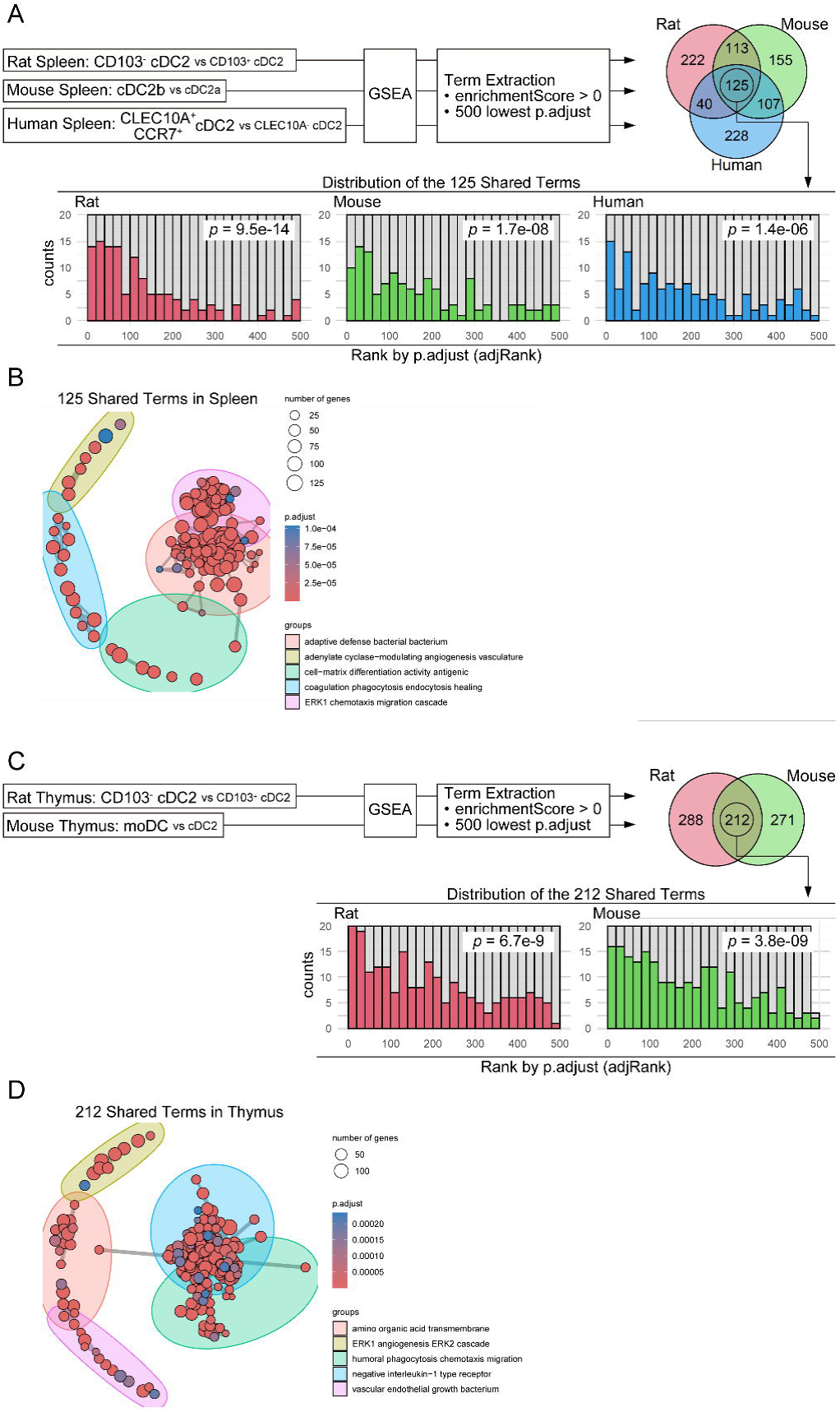
Commonly enriched terms across rat, mouse, and human CD103^-^ cDC2 equivalent subsets. (A, C) Comparison of enriched terms in splenic (A) and thymic (C) CD103-cDC2 or their equivalent subsets across species. GSEAs were performed as indicated in the figures. Skewed distributions of the *adjRank* within the shared terms were assessed using Wilcoxon rank-sum tests, and the resulting p-values are shown in the figure. (B, D) Enrichment maps of the 125 shared enriched terms in the spleen (B) and 212 shared enriched terms in the thymus (D).

For the splenic CD103^-^ cDC2 equivalents, 125 terms were found to be shared across the rat, mouse, and human datasets (Fig. 7A). Importantly, these shared terms exhibited significantly lower p.adjust values in all three species, suggesting stronger enrichment. To summarize the biological significance of these terms, an enrichment map was generated (Fig. 7B). The majority of the 125 common terms clustered into groups such as “adaptive defense bacterial bacterium” and “ERK1 chemotaxis migration cascade”, indicating enhanced roles in adaptive immunity.

Similarly, GSEA was performed for thymic CD103^-^ cDC2 equivalents in the mouse dataset. This analysis revealed 212 shared terms between rat and mouse thymic subsets (Fig. 7C). These common terms also showed lower p.adjust values, emphasizing their importance. The 212 shared terms were categorized into functional groups such as “negative interleukin-1 type receptor” and “humoral phagocytosis chemotaxis migration”, further indicating enhanced immunological activity of CD103^-^ cDC2 equivalents compared to CD103^+^ cDC2 equivalents (Fig. 7D).

The full lists of shared GO terms for splenic and thymic subsets are provided in supplementary tables 6 and 7 included in the *supplementary data and tables.xlsx*, respectively.

## Discussion

In this report, we discovered the rat CD103^-^ DC subsets in the thymus and spleen. Both flow cytometry and transcriptome analyses revealed that these CD103^-^ DC cells are novel subsets of CD172a^+^ cDC2. Moreover, while the classic CD103^+^ cDC2 subsets corresponded exclusively to cDC2a subpopulations, the novel rat CD103^-^ cDC2 subsets contained the counterparts of mouse and human cDC2 subpopulations: cDC2a, cDC2b, inf-cDC2, and moDC. Additionally, the splenic rat CD103^-^ cDC2 subset demonstrated enhanced immunogenic activity *in vitro*, consistent with the *in silico* analysis and previous findings in mouse cDC2b/moDC/inf-cDC2 (discussed later). Comparisons with mouse and human cDC2b and moDC subsets revealed that the shared terms tend to have lower p.adjust values and are associated with adaptive immunity. This suggests that these associated terms were not detected by chance, indicating that many of the related genes are strongly expressed and that such expression is conserved across species.

Regarding splenic cDC subsets, utilizing *Tbx21*^RFP-Cre^ mice, Brown *et al.* characterized and designated T-bet^+^ and T-bet^-^ cDC2s as cDC2a and cDC2b, respectively (40). They also identified counterparts of these cDC2a and cDC2b subsets in human spleens. The cDC2a subset is marked by high *Esam* expression, while the cDC2b subset expresses *Clec10a, Clec12a, Csf1r, Cx3cr1, Ccr2, Tlr8,* and *Tlr9*. Characteristics of the cDC2b subset align with the previously reported *Notch2*-independent cDC2s (41) and *KLF4*- dependent cDC2s (42,43). Although *Mgl2* is a marker for non-mDC cDC2 (similar to cDC2a in peripheral tissues) in the thymus (39), it is expressed on *KLF4*-dependent cDC2s (presumably cDC2b) in the lung and liver, but not in lymph nodes (43), suggesting that *Mgl2* could serve as another marker for peripheral cDC2b.

Despite observing the unique expression of many of the cDC2b/moDC signature genes (Fig. 5F, H), we did not find significant differences in *Tbx21* and *Esam* expression between the rat CD103^-^ cDC2 and CD103^+^ cDC2 subsets (Fig. 2E), suggesting either species differences or the need for more detailed gating. Additionally, both Klf4 and Notch2 were highly expressed in both subsets, indicating that while they are essential for differentiating cDC2a and cDC2b, their expression levels alone are insufficient as markers. At this point, CD103 can serve as a reliable marker to distinguish cDC2 subpopulations in rats. Utilizing CD103 and established antibodies, we can identify the rat cDC2 subsets that include cDC2b, inf-cDC2, and moDC counterparts.

Functionally, the cDC2b subset exhibits enhanced secretion of TNF-α, IL-6, and IL-10 upon TLR agonist or CpG stimulation (40,42). T cells primed by cDC2b produce higher levels of IFN-γ and IL-17 and show increased proliferation. DC-specific *Klf4* knockout mice have fewer Th17 cells in skin and skin-draining lymph nodes (42). Our rat splenic CD103^-^ cDC2s showed enrichment of TNF-α and IL-6-associated gene sets in *in silico* analyses (Fig. 3A and supplementary table 1) and induced greater T cell proliferation in MLR assays (Fig. 4A). These findings are consistent with known functional attributes of cDC2b.

However, the high potency of cDC2b does not imply that peripheral cDC2a subsets are low-potency cells. Notch2-dependent DCs (presumably cDC2a) are crucial for responses against xenogeneic red blood cells, heat-killed *Listeria monocytogenes* (44), and *Citrobacter rodentium* (45). These functions are likely mediated through antigen presentation via MHCII molecules. Since we could not assess the antigen-presenting ability of rat DC subsets due to the lack of appropriate TCR transgenic animals, it remains possible that the rat CD103^+^ cDC2 subsets play significant roles in adaptive immunity.

Inf-cDC2 is a relatively newly defined subpopulation, typically induced under inflammatory conditions (12,46). To our knowledge, no reports indicate the presence of this subset in healthy tissue, making the predominant expression of inf-cDC2 markers in the rat CD103^-^ cDC2 subsets from healthy animals unexpected (Fig. 5G). Further analysis, such as scRNA-seq or fate mapping, is required to determine whether inf-cDC2 counterparts exist in healthy rat organs. Researchers should consider this possibility when studying rat DCs.

As mentioned in the introduction, distinguishing cDC2b from moDC is challenging. One study suggested that, in a steady state, there are no *Zbtb46*^+^ DCs with a history of *Mafb* expression in the spleen (9). However, our splenic CD103^-^ cDC2 transcriptome exhibited significant expression of *Zbtb46* and *Mafb*, as well as other moDC markers such as *Cd14*, *Adgre1*, and *Csf1r* (Fig. 2F), suggesting that in rat spleens, some moDCs may be present even in a steady state.

Of the reported subsets of thymic cDC2 thus far, CD14^+^ moDCs were first described by Vobořil *et al*. in 2020 (39). These moDC cells predominantly express monocyte/macrophage gene signature molecules and some chemokine receptors such as *Cxcr2, Ccr1, Ccr5, Cxcr4, Cx3cr1,* and *Ccr2*. It has been shown that chemokines secreted from CpG-stimulated TECs recruit moDCs to the thymus, ultimately facilitating antigen transfer from TECs to moDCs. Therefore, TEC-specific disruption of TLR signaling leads to reduced moDC recruitment, impaired Treg development, and failure to prevent experimental colitis. Additionally, moDCs are reported to incorporate TEC-derived antigens more effectively than other cDC subpopulations, highlighting their essential role as antigen-presenting cells in the negative selection process (47).

Although the term moDC was not used to describe them, CX3CR1^+^ cDC2s have been reported to localize trans-endothelially in the thymus, capturing blood-borne molecules (5), and play a critical role in inducing gut commensal bacteria-specific T cells (6).

Our rat thymic CD103^-^ cDC2 subset showed a high similarity score to the mouse thymic moDC subset (Fig. 5C), and mouse moDC markers were predominantly expressed in the rat thymic CD103^-^ cDC2 subset (Fig. 5D, E). These results suggest that the rat thymic CD103^-^ cDC2 subset is either the counterpart of the mouse moDC subset or includes it. GSEA results further revealed that both the rat CD103^-^ cDC2 (Fig. 3C) and mouse moDCs (Fig. 7C, D) are enriched with various immune activation-associated terms.

However, non-moDC cDC2 cells are also crucial in the negative selection. Breed *et al.* found that the CD301b^+^ (*Mgl2*, the non-moDC cDC2 marker) cDC2 subset exhibits enhanced MHCII expression, H2-DM/H2-DO ratio, and I-A^b^ CLIP expression, suggesting improved antigen presentation (33). Indeed, CD301b^+^ cell-eliminated mice display a broader TCR repertoire, implying impaired negative selection.

In line with the reported roles of both moDC and non-moDC cDC2 subsets in negative selection, our RTOC experiments demonstrated that both the rat CD103^-^ and CD103^+^ cDC2 subsets suppressed thymocyte generation and shifted their differentiation toward CD8SP, indicating enhanced negative selection (Fig. 4B). It is difficult to conclude definitively whether one subset is more significant in this process; their importance likely varies depending on experimental context and antigen type. Notably, only the CD103^+^ cDC2 subset promoted the maturation of the generated CD4SP thymocytes (Fig. 4D), suggesting that negative selection and subsequent maturation may be regulated by different cell populations and molecules.

In contrast to the immune-associated terms enriched in the CD103^-^ cDC2 subsets, the p.adjust values of cell cycle-associated terms enriched in the rat CD103^+^ cDC2 were high (Fig. 3A, C), and their absolute values for average fold changes and differential scores were low (Fig. 3E, F). Additionally, we did not observe significant differences in proliferative marker Ki-67 expression between the rat CD103^-^ cDC2 and the rat CD103^+^ cDC2 subsets by flow cytometry (data not shown). Supporting these findings, mitotic DCs are a minor component of mouse and human cDC2 subsets (Fig. S5A, B, S6A). Therefore, it is noteworthy that immune-associated terms in the rat CD103^+^ cDC2 subsets were even less significant than these weak cell cycle-associated terms. This may imply that the immunological activity of the rat CD103^+^ cDC2 subsets, if any, requires a priming event(s) to become fully apparent.

In conclusion, we demonstrated the presence of novel CD103^-^ conventional DC subsets in the rat spleen and thymus. These are subpopulations of cDC2 and include counterparts of mouse and human cDC2a, cDC2b, inf-DC2, and moDC subpopulations, while the classic rat CD103^+^ cDC2 subsets correspond exclusively to cDC2a subpopulations. Additionally, the expression of adaptive immunity-associated terms was conserved across rat, mouse, and human CD103-cDC2 and their equivalents. Although identifying cDC2 subpopulations in mice and humans is challenging, rat researchers can reliably identify these potent subsets using well-established antibodies.

## Author Contributions

Yasushi Sawanobori developed the concept, designed and performed all experiments and analyses, and wrote the codes and manuscript. Tadayuki Ogawa, Yusuke Kitazawa, Hisashi Ueta, and Nobuko Tokuda provided the research environment, contributed materials, and supervised the research.

## Acknowledgments

The authors would like to thank Ms. Yasuko Nonaka for her expert operation of the cell sorter. We are deeply grateful to Prof. Emer. Kenjiro Matsuno, for his insightful comment years ago: “There would be CD103^-^ DCs in rat tissues.” Lastly, we thank ChatGPT, who greatly supports me in learning R and English writing.

## Figure legends

**Supplementary Figure 1.**
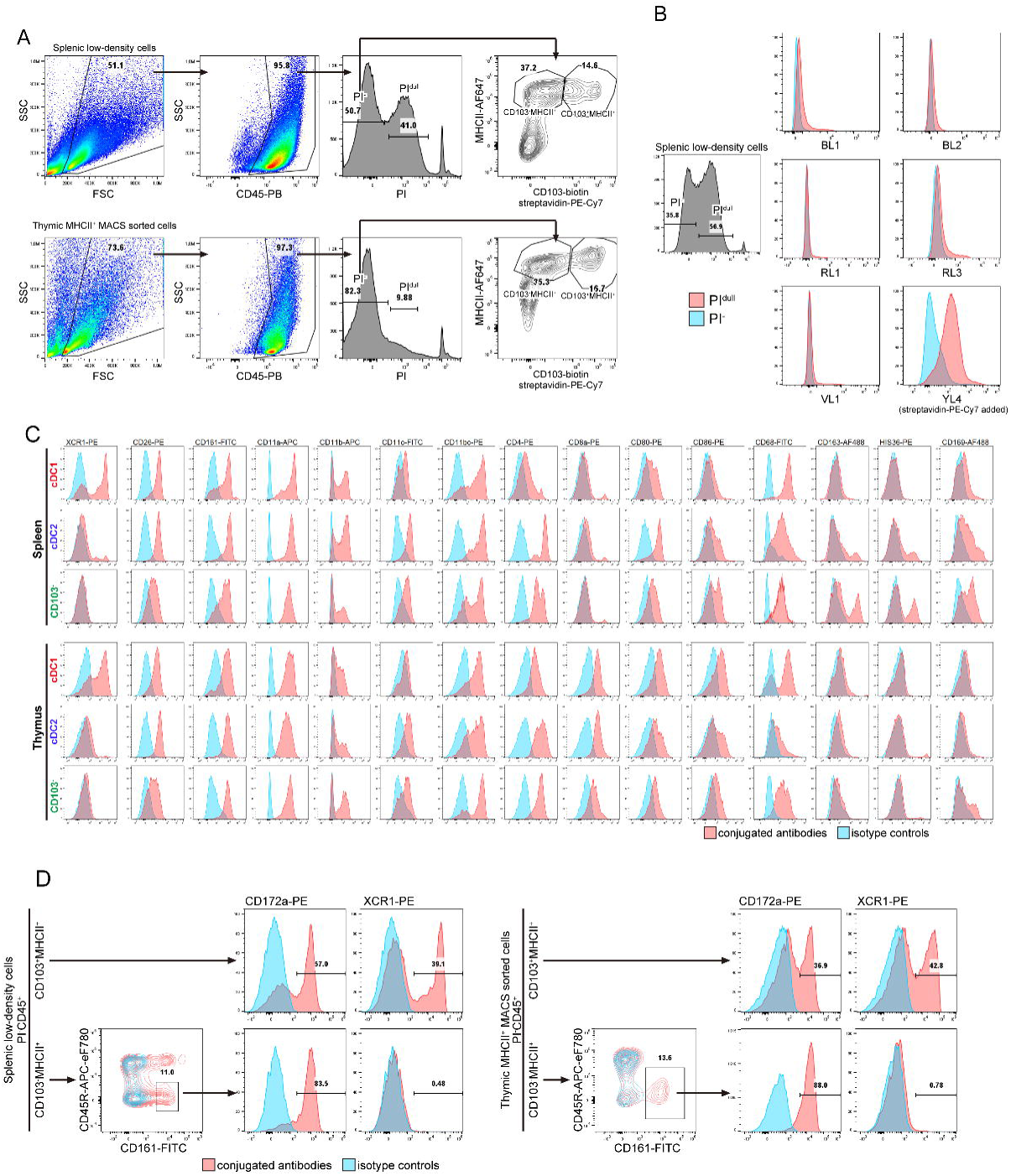
Gating strategy for rat DC analysis and surface molecule expression of rat DC subsets. (A) Gating strategy for the three rat DC subsets in the spleen and thymus. (B) Autofluorescence of splenic PI^dull^ cells. (C) Surface molecule expression of the three DC subsets in the thymus and spleen. (D) CD172a and XCR1 expression on CD103^+^MHCII^+^ cDC subsets and the CD103^-^ MHCII^+^CD45R^-^CD161^+^ novel cDC subset in the thymus and spleen. All data were collected from 8-week-old rats.

**Supplementary Figure 2.**
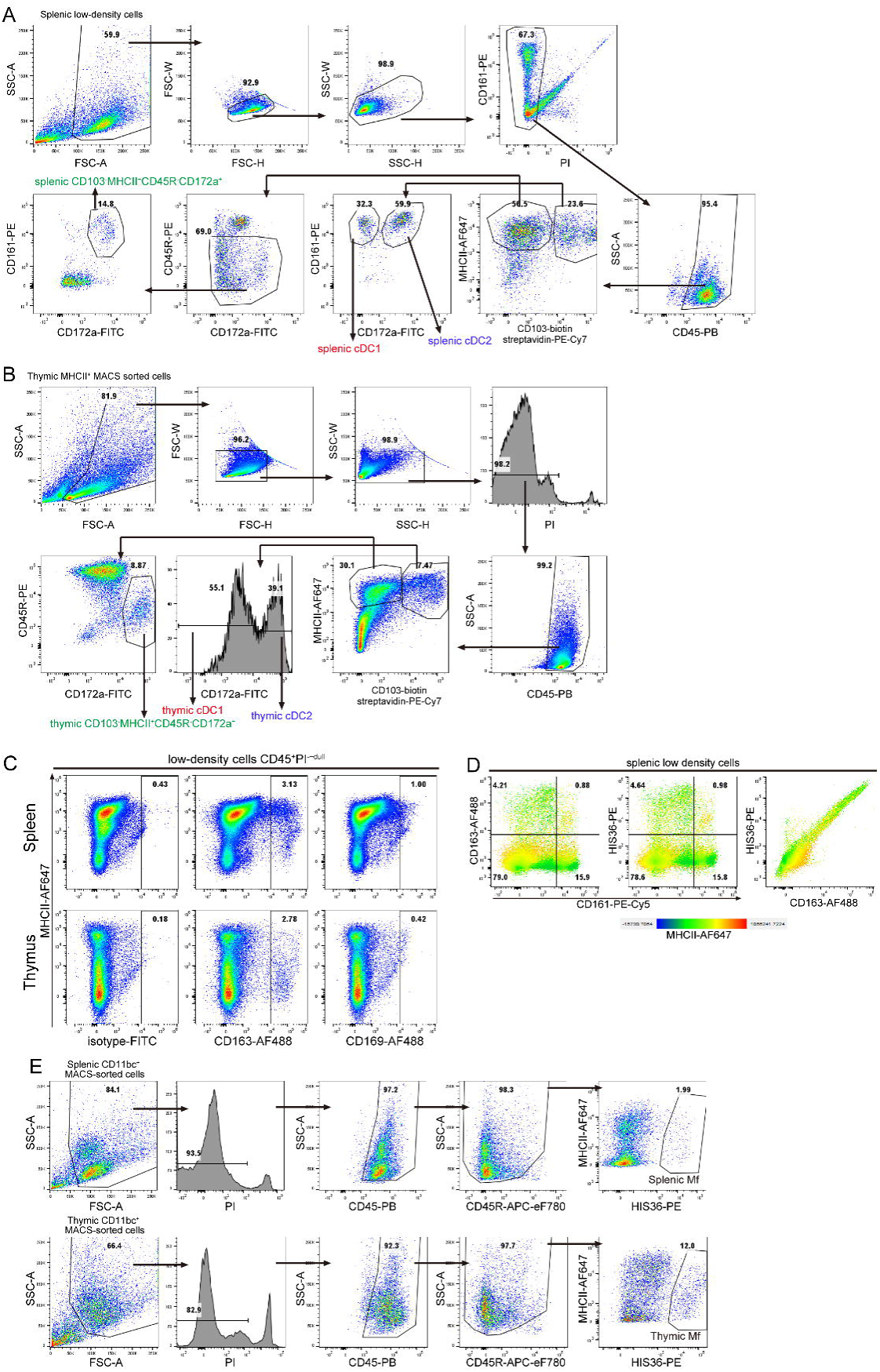
Gating strategies for cell sorting. (A, B) Gating strategies for cell sorting of splenic DCs (A) and thymic DCs (B). (C) Expression of macrophage markers in low-density cells. (D) Identical staining with anti-CD163 and HIS36 antibodies. (E) Gating strategies for sorting of splenic and thymic macrophages. Rats were 10 weeks old (A, B), 12 weeks old (C), 8 weeks old (D), and 9 weeks old (E).

**Supplementary Figure 3.**
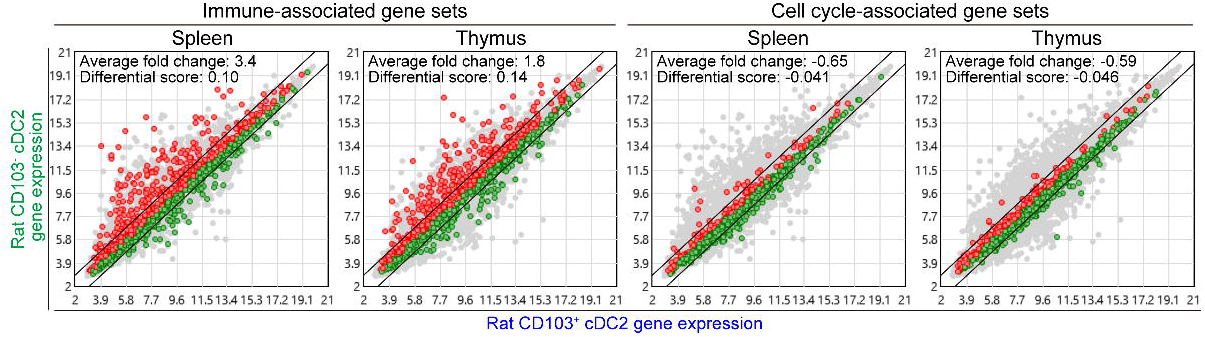
Immune-associated and cell cycle-associated gene sets applied to scatter plots of rat transcriptomes. Gene sets for immune-associated (GO:0006909, GO:0002253, GO:0001819) and cell cycle-associated (GO:0006260, GO:0007059, GO:0044839) terms were extracted and applied to scatter plots of rat CD103^-^ and CD103^+^ cDC2 transcriptomes.

**Supplementary Figure 4.**
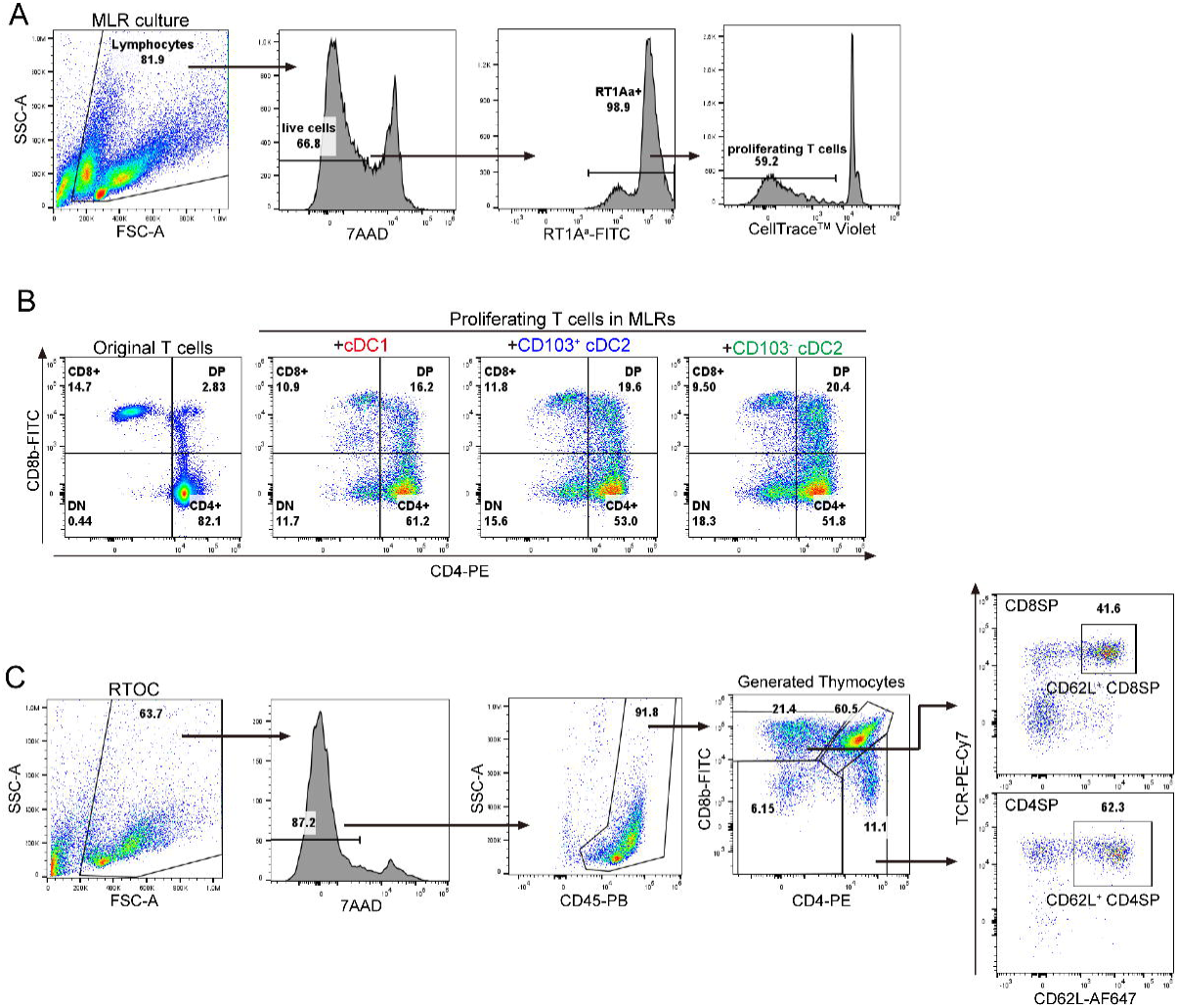
Gating strategies and representative results of MLR and RTOC analyses. (A) Gating strategy and a representative result of the MLR, showing an excerpt from MLRs with 1000 CD103^-^ cDC2 cells. (B) CD4−CD8 profiles of T cells in MLR experiments. T cells were stimulated by 1000 DCs. (C) Gating strategy and a representative result of the RTOC, showing an excerpt from no DC-RTOCs.

**Supplementary Figure 5.**
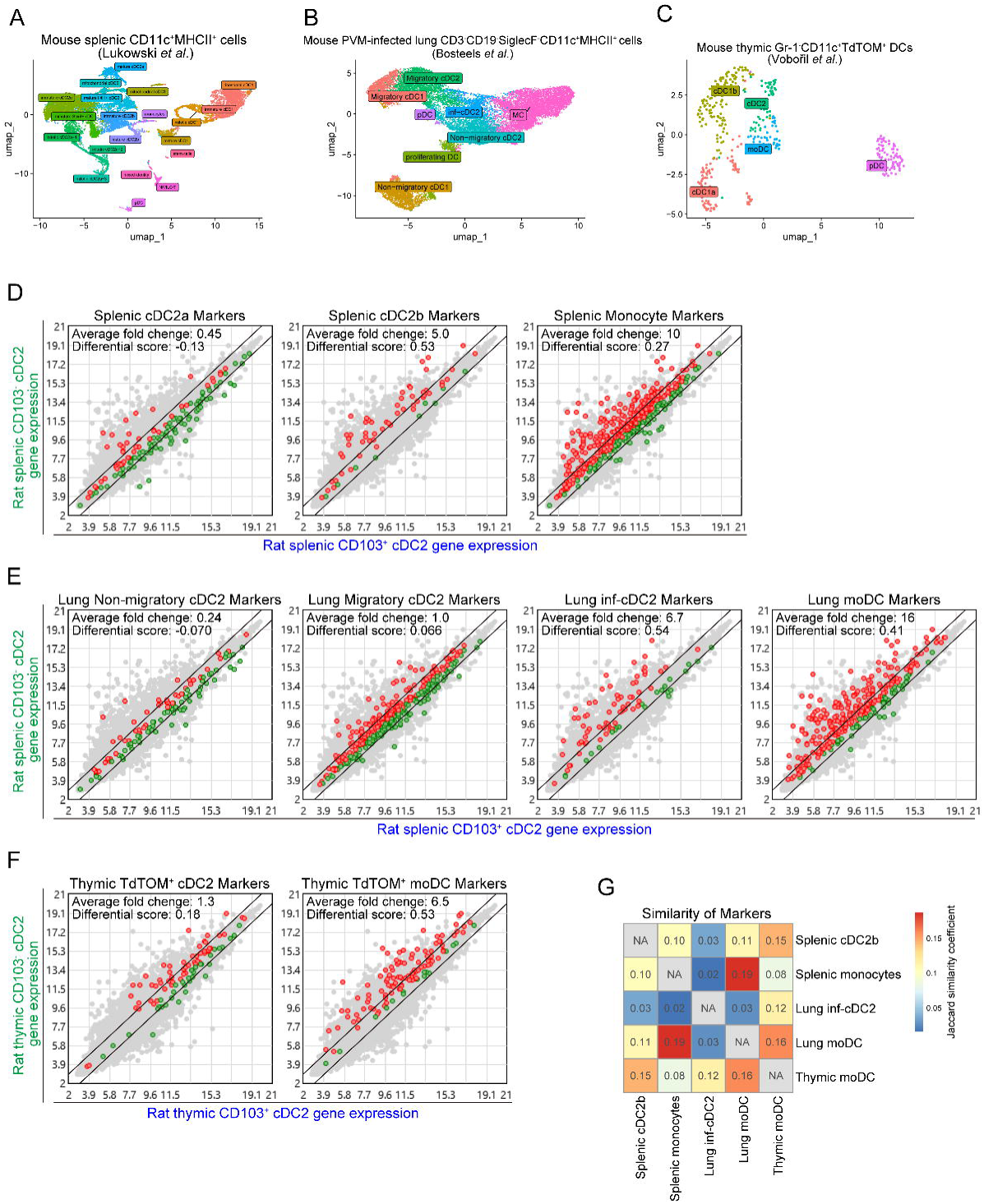
2D projections of mouse scRNA-seq and mouse cDC2 subpopulation markers applied to scatter plots of rat transcriptomes. (A–C) Umap projections of mouse splenic DC (A), PVM-infected lung DC (B), and thymic TdTOMATO-incorporating DC (C) subsets. For (A), clusters were taken from the original metadata, but the projection was newly generated. For (B), both the clustering and projection were taken from the original metadata. For (C), both the clustering and projection were newly generated to reproduce the original research. (D–F) Gene marker sets from mouse splenic (D), lung (E), and thymic (F) cDC2 subpopulations were applied to the rat microarray data. Average fold changes and differential scores were then calculated. (G) Jaccard similarity coefficients calculated between the mouse marker gene sets displayed as a heatmap.

**Supplementary Figure 6.**
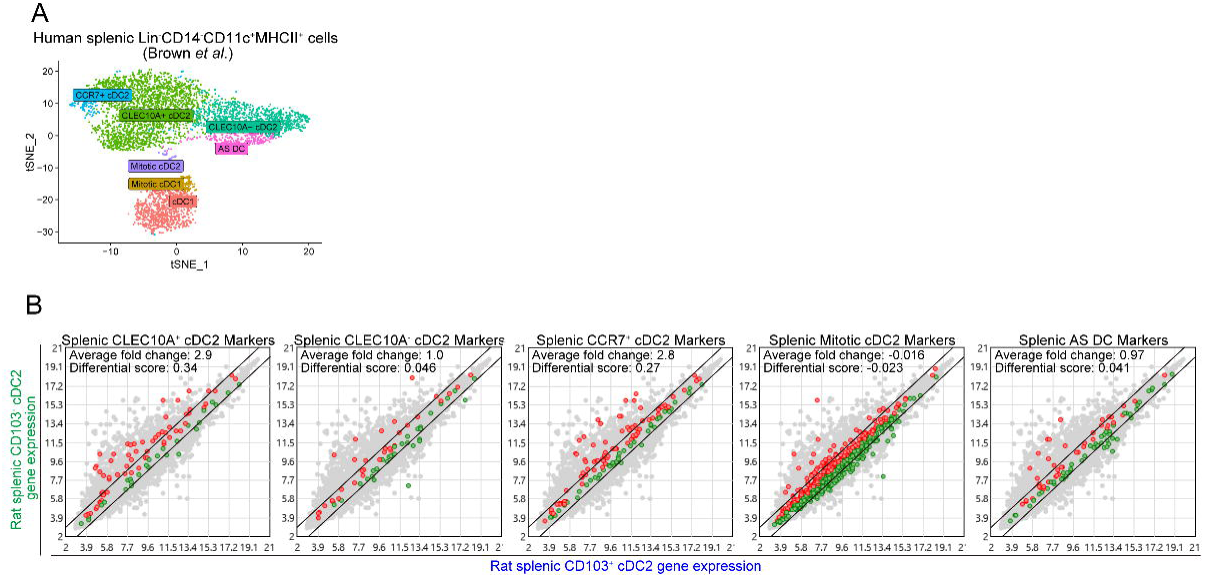
2D projection of human scRNA-seq and human cDC2 subpopulation markers applied to rat transcriptomes. (A) tSNE projections of human splenic DC subpopulations. The clustering and projection were taken from the original metadata. (B) Gene marker sets from human splenic cDC2 subpopulations were applied to the rat microarray data. Then, average fold changes and differential scores were calculated.

